# Significant cross-species gene flow detected in the *Tamias quadrivittatus* group of North American chipmunks

**DOI:** 10.1101/2021.12.07.471567

**Authors:** Jiayi Ji, Donavan J. Jackson, Adam D. Leaché, Ziheng Yang

## Abstract

In the past two decades genomic data have been widely used to detect historical gene flow between species in a variety of plants and animals. The *Tamias quadrivittatus* group of North America chipmunks, which originated through a series of rapid speciation events, are known to undergo massive amounts of mitochondrial introgression. Yet in a recent analysis of targeted nuclear loci from the group, no evidence for cross-species introgression was detected, indicating widespread cytonuclear discordance. The study used heuristic methods that analyze summaries of the multilocus sequence data to detect gene flow, which may suffer from low power. Here we use the full likelihood method implemented in the Bayesian program BPP to reanalyze these data. We take a stepwise approach to constructing an introgression model by adding introgression events onto a well-supported binary species tree. The analysis detected robust evidence for multiple ancient introgression events affecting the nuclear genome, with introgression probabilities reaching 65%. We estimate population parameters and highlight the fact that species divergence times may be seriously underestimated if ancient cross-species gene flow is ignored in the analysis. Our analyses highlight the importance of using adequate statistical methods to reach reliable biological conclusions concerning cross-species gene flow.

## INTRODUCTION

Genomic sequence data are a rich source of information concerning the history of species divergences and cross-species gene flow. The past two decades have seen widespread use of genomic data to infer hybridization or introgression (Mallet *et al*., 2016). Cross-species hybridization/introgression has been detected in a variety of species including Arabidopsis (Arnold *et al*., 2016), butterflies (Martin *et al*., 2013), corals (Mao *et al*., 2018), lizards (Finger *et al*., 2021), birds (Ellegren *et al*., 2012), mammals (Kumar *et al*., 2017; Chan *et al*., 2013; Shi and Yang, 2018), and hominins (Nielsen *et al*., 2017). The studies have considerably enriched our understanding of the evolutionary dynamics of introgressed genes, and the role of introgression in speciation and ecological adaptation (Payseur and Rieseberg, 2016; Martin and Jiggins, 2017).

A number of statistical methods have also been developed to analyze genomic sequence data to detect gene flow between species and to estimate its strength (as measured by the introgression probability or migration rate). Heuristic methods make use of summary statistics and include the popular *D*-statistic or ABBA-BABA test (Patterson *et al*., 2012), which uses the site-pattern counts for a species quartet to test for the presence of gene flow between two non-sister species, SNaQ (Solis-Lemus and Ane, 2016), and HyDe (Blischak *et al*., 2018). Full likelihood methods use the multilocus sequence alignments directly and include the Bayesian implementations of the introgression model in PhyloNet/MCMC-SEQ (Wen and Nakhleh, 2018), *BEAST (Zhang *et al*., 2018), and BPP (Flouri *et al*., 2020), as well as the MCMC implementation of the continuous-migration model (also known as the isolation-with-migration or IM model) of Hey *et al*. (2018). See Jiao *et al*. (2021) for a recent review. In theory, full-likelihood methods may be expected to be more powerful because they make an efficient use of information in the data. However, summary and likelihood methods for inferring cross-species gene flow are seldom applied to the same real datasets with their utilities evaluated, partly because likelihood methods typically involve intensive computation and may not be computationally feasible for genome-scale datasets. In this regard, it is noteworthy that the BPP implementation of the MSci model has been successfully applied to genomic datasets of more than 10,000 loci (Flouri *et al*., 2020, table 1; Thawornwattana *et al*., 2021, table S4).

**Table 1.** Summary of evidence for mitochondrial introgression in the *T. quadrivittatus* group (Sullivan *et al*., 2014)

| Species | Region | Distribution | Introgression | Source |
| --- | --- | --- | --- | --- |
| *T. bulleri* | M | Allopatric | No |  |
| *T. canipes* (C) | GB/RM | Allopatric | No |  |
| *T. cinereicollis* (I) | GB/RM | Parapatric | Yes | Not assignable |
| *T. dorsalis* (D) | GB/RM | Parapatric | Yes | C/U/Q/Not assignable |
| *T. durangae* | M | Allopatric | No |  |
| *T. palmeri* | GB/RM | Allopatric | Untested |  |
| *T. quadrivittatus* (Q) | GB/RM | Parapatric | Yes | Not assignable |
| *T. rufus* (R) | GB/RM | Allopatric | No |  |
| *T. umbrinus* (U) | GB/RM | Parapatric | Yes | Not assignable |

Note.— Geographic regions include Great Basin (GB), Rocky Mountains (RM), and Mexico (M). Single letter codes are for the six species included in the nuclear data analysis.

The *Tamias* chipmunks are a diverse group of at least 23 distinct species, occupying a variety of habitats in the western United States (Patterson and Norris, 2016). Recent molecular phylogenetic work on this group has elucidated a complex history of radiative speciations and cross-species gene flow involving morphologically and ecologically diverse lineages (Good and Sullivan, 2001; Good *et al*., 2003). The *Tamias quadrivittatus* group of chipmunks consists of about 9 species that are currently distributed across the Great Basin along with the central and southern Rocky Mountains in North America (fig. 1). Previous work on *Tamias* has highlighted the importance of genital morphology, specifically the baculum (a bone found in the penis) in male chipmunks, as a reliable indicator of species limits (Patterson and Thaeler Jr, 1982; White, 2010). The biogeographic history of the group likely included large range fluctuations that have periodically resulted in isolation and secondary contact among species, which would have provided opportunities for hybridization and/or introgression (Good *et al*., 2003). The current distributions of species in the group has extensive regions of overlap and broad parapatry in ecological transition zones (fig. 1), with instances of both allopatry and parapatry, and the determinants of current distributions are thought to be related primarily to competitive exclusion and ecological preference (Brown, 1971; Heller, 1971; Root *et al*., 2001). This system provides an intriguing opportunity to investigate the effects of introgression on genetic variation within and among species.

**Figure 1:**
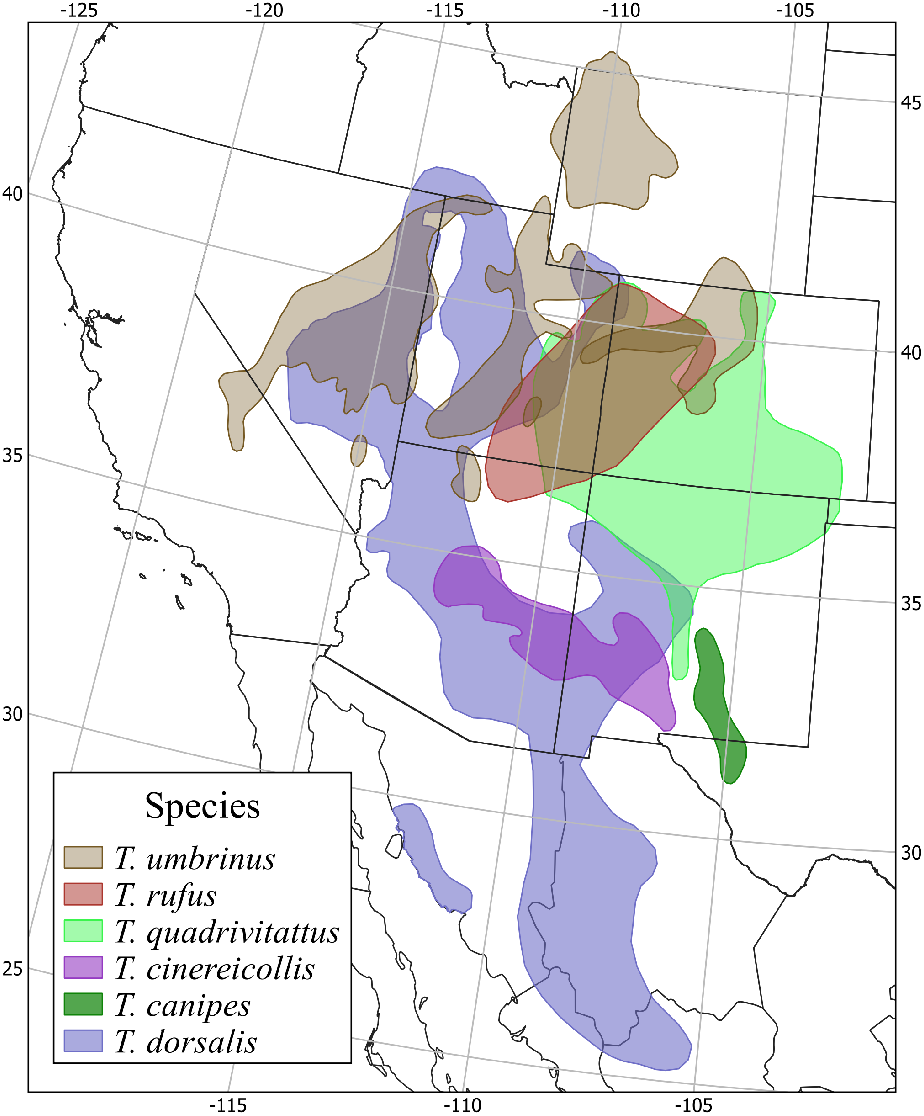
Geographic distributions of the six chipmunk species in the *Tamias quadrivittatus* group, based on data downloaded from the IUCN (https://www.iucnredlist.org/).

Hybridization between chipmunk species was reported in early studies (Good and Sullivan, 2001; Good *et al*., 2003, 2008; Hird *et al*., 2010). Work in the past decade has documented widespread mitochondrial introgression among species of the group (Reid *et al*., 2012; Sullivan *et al*., 2014; Sarver *et al*., 2017, 2021), often characterized by asymmetric mitochondrial introgression, possibly due to bacular morphology, that has been identified in at least 6 species (Good *et al*., 2003, 2008; Reid *et al*., 2012; Sullivan *et al*., 2014). Recent work on six species in the *T. quadrivittatus* group found that four of them exhibited clear evidence of introgressed mitochondrial DNA: *T. cinereicollis, T. dorsalis, T. quadrivittatus*, and *T. umbrinus* (table 1). The cliff chipmunk (*T. dorsalis*) was involved in local introgression with multiple other species, receiving mtDNA from whichever congener chipmunk it came into contact with. However, populations of *T. dorsalis* that are geographically isolated carry mtDNA haplotypes that are unique to the species (Sullivan *et al*., 2014; Sarver *et al*., 2017). This illustrates a recurrent trend of mitochondrial introgression exhibited in *Tamias*, that range overlap in transition zones plays an important role in gene flow (Brown, 1971; Bi *et al*., 2019).

Sarver *et al*. (2021) used a targeted sequence-capture approach to sequence thousands of nuclear loci (mostly genes or exons) to estimate the species phylogeny of the *T. quadrivittatus* group and to infer possible nuclear introgression. The program HyDe (Blischak *et al*., 2018) was used to infer gene flow. Surprisingly, no significant evidence for gene flow involving the nuclear genome was detected between any species in the group, despite the evidence for widespread mitochondrial introgression. We note that HyDe, like the *D*-statistic (Patterson *et al*., 2012), is a heuristic method that uses the four-taxon site-pattern counts pooled across the genome. It cannot detect gene flow between sister species at all, and, for nonsister species, fails to use information in the fluctuation in genealogical history across the genome caused by the stochastic process of coalescent and introgression (Zhu and Yang, 2021; Jiao *et al*., 2021); see Discussion for a characterization of the amount of information used by HyDe.

To test whether the lack of evidence for nuclear introgression found by Sarver *et al*. (2021) may be due to the inefficiency of analytical methods used, here we re-analyze the data of Sarver *et al*. (2021) using the BPP program (Flouri *et al*., 2018, 2020), which includes a full likelihood implementation of the multispeciescoalescent-with introgression (MSci) model. Borrowing ideas from stepwise regression or Bayesian variable selection, we add introgression events sequentially onto the binary species tree to construct a joint MSci model with multiple introgression events. Our analysis revealed robust evidence for multiple ancient introgression events, involving both sister species and nonsister species. We use computer simulation to verify that the lack of power of the summary method is responsible for the opposite conclusions reached in the two analyses. We then assess the impact of ignoring introgression on estimation of population parameters, highlighting serious biases in species divergence time estimation when introgression exists and is ignored. Our results highlight the power of coalescent-based full-likelihood methods in the analysis of genomic datasets to infer the history of species divergence and gene flow.

## MATERIALS AND METHODS

### Chipmunk data

The dataset, generated and analyzed by Sarver *et al*. (2021), includes 1060 nuclear loci from six chipmunk species: *T. rufus* (R), *T. canipes* (C), *T. cinereicollis* (I), *T. umbrinus* (U), *T. quadrivittatus* (Q) and *T. dorsalis* (D) (with 5, 5, 9, 10, 11, 11 individuals, respectively), as well as the outgroup *T. striatus* (3 individuals). We included all individuals whether or not their mtDNA was likely to be introgressed. Due to lack of a reference genome, Sarver *et al*. (2021) assembled genomic loci (targeted genes or exons) into contigs using an approach called Assembly by Reduced Complexity (ARC). Filters were then applied to remove missing data (contigs not present across all individuals) and sequences with likely assembly errors. The procedure generated a dataset of 1060 loci, with sequence length ranging from 14 to 1026 bp among loci and the number of variable sites from 0.33% to 15.2%.

High-quality heterozygous sites in the data, as identified by high mapping quality and depth of coverage, are represented using IUPAC ambiguity codes. They are accommodated using the analytical integration algorithm implemented in BPP (Flouri *et al*., 2018; Gronau *et al*., 2011). This takes the unphased genotype sequences as data and averages over all possible heterozygote phase resolutions, using their relative likelihoods based on the sequence alignment at the locus as weights (Huang *et al*., 2021).

### Species tree estimation

We estimated the species tree under the multispecies coalescent (MSC) model without gene flow implemented in BPP version 4 (Flouri *et al*., 2018; Rannala and Yang, 2017). This is the A01 analysis (`speciedemilitation=0`, `speciestree=1`) (Yang, 2015).

We assigned inverse-gamma (IG) priors to parameters in the MSC model: *θ* ∼ IG(3, 0.002) with mean 0.001 for population size parameters and *τ*_0_ ∼ IG(3, 0.01) with mean 0.005 for the age of the root. The shape parameter *α* = 3 means that those priors are diffuse, while the prior means are based on estimates from preliminary runs. Note that both *θ* s and *τ*s are measured in the expected number of mutations per site. The inverse-gamma is a conjugate prior for *θ* and allows the *θ* parameters to be integrated out analytically, leading to improved mixing of the Markov chain Monte Carlo (MCMC) algorithm. We conducted 10 replicate MCMC runs, using different starting species trees. Each run generated 2 × 10^5^ samples, with a sampling frequency of 2 iterations, after a burn-in of 16,000 iterations. Each run took about 70 hours using one thread on a server with Intel Xeon Gold 6154 3.0GHz processors. Convergence was confirmed by consistency between runs. All runs converged to the same species tree (fig. 2), with ∼ 100% posterior probability, which had the same topology as the tree inferred by Sarver *et al*. (2021).

**Figure 2:**
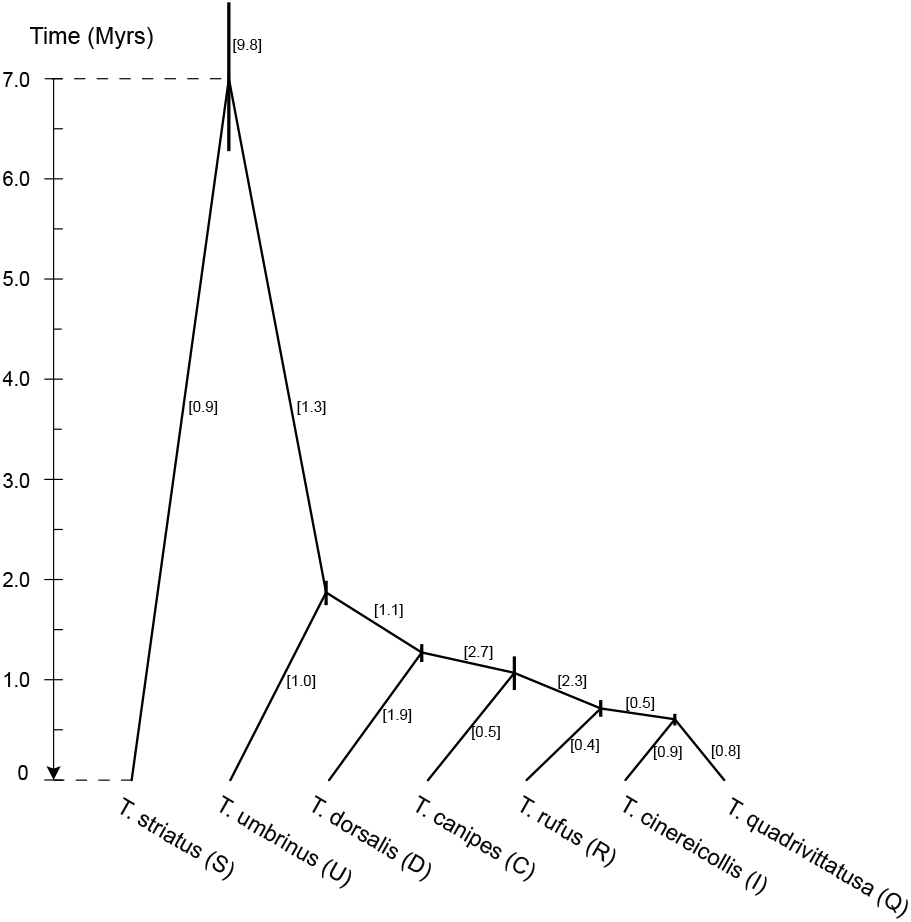
Species tree for the *T. quadrivittatus* group with *T. striatus* used as the outgroup. Branch lengths represent the posterior means of divergence times (*τ*s) estimated from BPP analysis of the full data of 1060 loci under the MSC model with no gene flow, with node bars indicating the 95% HPD intervals. Posterior means of population size parameters (*θ* s) are in brackets. A minimum divergence time of 7 Myrs for the outgroup *T. striatus* is used to convert the *τ* estimates into absolute times.

### Stepwise construction of the introgression model

As the species tree is well supported, apparently unaffected by small amounts of cross-species introgression, we used the species tree to build an introgression model with multiple introgression events. Our procedure is similar to stepwise regression, the step-by-step method for constructing a regression model that involves adding or removing explanatory variables based on a criterion such as an *F*-test or *t*-test.

Our procedure has two stages. In the first stage, we used BPP to fit a number of introgression models, each with only one introgression event, and rank candidate introgression events by their strength (indicated by the introgression probability *φ*). The analyses of Sarver *et al*. (2021) suggest that mitochondrial introgression affected mostly four species: *T. umbrinus* (U), *T. dorsalis* (D), *T. quadrivittatus* (Q) and *T. cinereicollis* (I) (Sarver *et al*., 2021). We considered introgression events involving all possible pairs among those four species, as well as another species, QI, the common ancestor of *T. cinereicollis* and *T. quadrivittatus* (fig. 2). The dataset of 1060 loci was analyzed under an MSci model involving only one introgression event, estimating the introgression probability (*φ*) and introgression time (*τ*). Two replicate runs were conducted for each analysis to confirm consistency between runs, and MCMC samples from the two runs were then combined to produce posterior estimates of parameters. This analysis provides a ranking of the introgression events by the introgression probability, and only those with the posterior mean 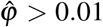 and with the 95% highest probability density (HPD) credibility interval (CI) excluding 0.001 considered further (Huang *et al*., 2021). This criterion is somewhat arbitrary and is used with the presupposition that an introgression event will be included in the model only if there is strong evidence for it.

In the second stage, we added introgression events onto the binary species tree (fig. 2) sequentially in the order of decreasing strength (introgression probability). To reduce the computational cost and to check for robustness of the analysis, this stepwise procedure was applied to two subsets of the 1060 loci: the first half and the second half, each of 530 loci. The priors used were as above. With multiple introgression events in the model, longer MCMC runs were needed. We thus extended the MCMC runs to be *k*-times as long if the model involved *k* introgression events. Three replicate runs were performed to check consistency between runs. Samples from the replicate runs were then combined to produce posterior summaries. At each step, the added introgression event was retained if it met the same cutoff as above in either of the two data subsets.

Our procedure produced a joint introgression model with five unidirectional introgression events and one bidirectional introgression event. The joint model was then applied to the full dataset of 1060 loci to estimate the population parameters including introgression probabilities, introgression times, species divergence times, and population sizes (fig. 3), using the same prior settings. We conducted 3 replicate runs, using a burn-in of 50,000 iterations and then taking 2 × 10^6^ samples, sampling every 2 iterations. Each run took 400 hrs.

**Figure 3:**
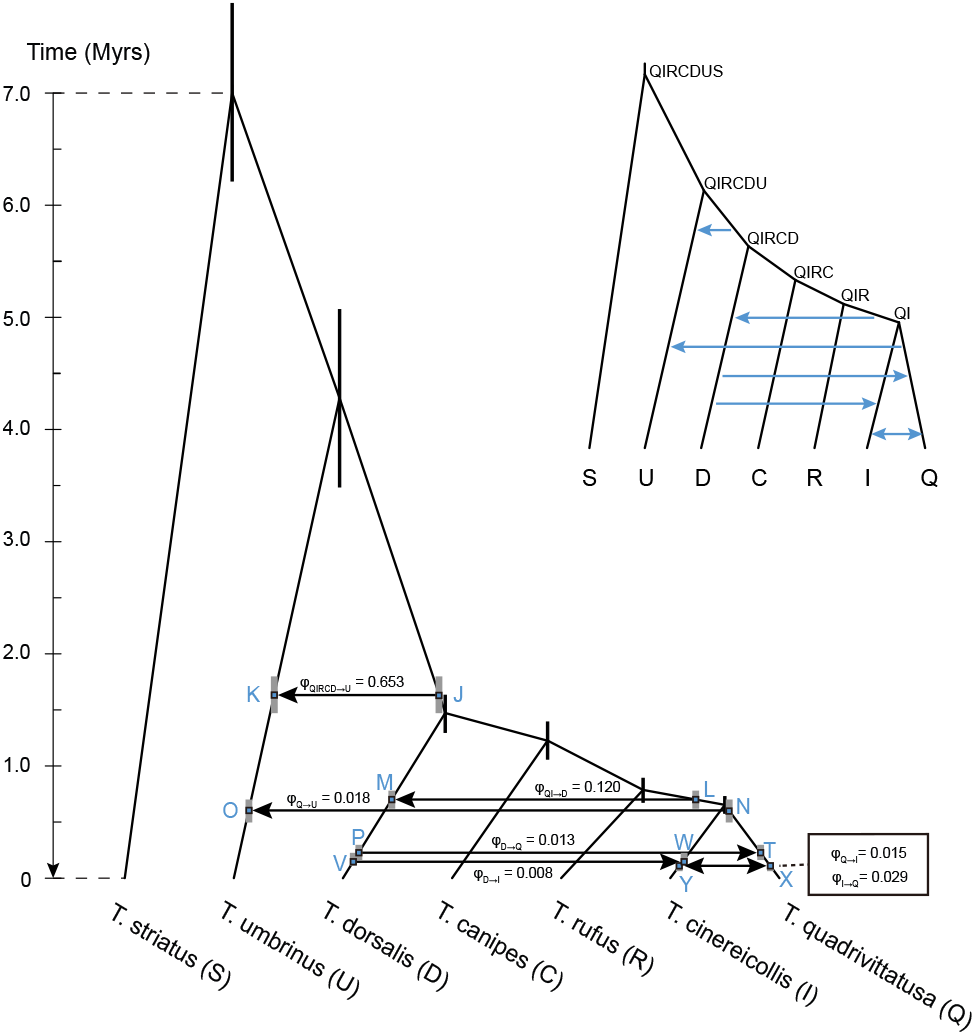
The joint introgression model constructed in this study with five unidirectional and one bidirectional introgression events showing parameter estimates from BPP analysis of the full data of 1060 loci. Branch lengths and node bars represent posterior means and 95% HPD intervals of divergence/introgression times (*τ*s). Nodes created by introgression events are labeled, with the labels used to identify parameters in table 3. The MSci model includes 6 species divergence times and 6 introgression times (*τ*s), 25 population size parameters (*θ* s), and 7 introgression probabilities (*φ*s).

### Simulation to evaluate the power of heuristic and likelihood methods to infer gene flow

Our BPP analyses of the nuclear genomic data revealed strong evidence for multiple introgression events (fig. 3), whereas no evidence for gene flow was detected by Sarver *et al*. (2021) using HyDe to analyze the same data. To explore whether the opposing results may be explained by the different powers of the methods used, we conducted two sets of simulations.

In the first set, we aimed to determine the power of HyDe and BPP to detect a strong introgression event using data similar to the *Tamias* dataset but at different numbers of loci. We simulated replicate datasets using the parameter estimates obtained under the joint model of figure 3 from the full data (see table 3 below). We simulated four (phased) sequences per species per locus, with the sequence length to be 200 sites. The number of loci was *L* = 100, 200, 400, 800, and 1600. The number of replicates is 100. The strongest introgression from our analysis is from QIRCD→U, between sister species. Since HyDe cannot detect introgression between sister species, we examined the next introgression event, from QI→D. We analyzed the data for the quartet of species, Q, R, D and S (outgroup), using BPP (Flouri *et al*., 2018, 2020) and Hyde (Blischak *et al*., 2018) to detect introgression from Q→D. In the HyDe test, species Q was treated as the ‘hybrid’ while R and D as the two parents (*P*_1_ and *P*_2_) (Blischak *et al*., 2018).

In the second set of simulations, we tested whether BPP achieves good accuracy and precision for parameter estimation from datasets of this size (e.g., six species, 1,000+ loci). We simulated two replicate datasets of 1060 loci using the parameter estimates (posterior means) from the full data (fig. 3, table 3). Each dataset had the same size as the original, i.e., 5, 5, 9, 10, 11, 11, 3 unphased sequences per locus, respectively, for species R, C, I, U, Q, D and S, with the sequence length to be 200 sites. The simulated datasets were analyzed using BPP under the joint MSci model (fig. 3) to estimate all parameters, for comparison with the empirical estimates (which were the true parameter values for the simulated data).

## RESULTS

### Species tree estimation

We analyzed the full data of 1060 loci under the MSC model without gene flow to estimate the species tree. The ten replicate runs using different starting species trees converged to the same maximum *a posteriori* probability (MAP) tree, with posterior probability ∼ 100% (fig. 2). Sarver *et al*. (2021) recovered the same species tree topology in their analysis of the same data using ASTRAL (Mirarab and Warnow, 2015) and SVDqu-ARTETS (Chifman and Kubatko, 2014), although with weaker support for some nodes, e.g., concerning the placement of *T. rufus*. The differences in support may be due to the fact that ASTRAL and SVDquartets use summaries of the multilocus sequence data that are not sufficient statistics, and are thus less efficient than the full likelihood method implemented in BPP (Xu and Yang, 2016; Zhu and Yang, 2021).

### Stepwise construction of the introgression model

In the first stage of our procedure, we fitted introgression models, each involving one introgression event, using full dataset of 1060 loci. We considered introgression events between each pair of the five species: *T. cinereicollis* (I), *T. dorsalis* (D), *T. quadrivittatus* (Q), and *T. umbrinus* (U), and the ancestral species QI (fig. 2). Introgression events that passed our cutoffs are listed in table 2. Introgression from QI into D had the highest probability, *>* 10%, while six more events had *φ >* 5%: Q→D, D→QI, QI→U, I→D, Q→I, and I→Q. We note that the introgressions between Q and I, and between QI and D, were significant in both directions and the estimated introgressions times were close (table 2). We thus replaced the two unidirectional introgression events by one bidirectional introgression in further analyses (model D in Flouri *et al*., 2020).

**Table 2.** Posterior means and 95% HPD CIs (in parentheses) for introgression probability (*φ*) and introgression time (*τ*) in the separate introgression analysis.

| Introgression | $\phi$ | $\tau (\times 10^{-3})$ |
| --- | --- | --- |
| QIRCD → U | 0.6215 (0.3907, 0.8243) | 0.896 (0.784, 1.004) |
| QI → D | 0.1187 (0.0866, 0.1499) | 0.337 (0.311, 0.367) |
| Q → D | 0.0779 (0.0509, 0.1026) | 0.297 (0.253, 0.328) |
| D → QI | 0.0707 (0.0384, 0.1058) | 0.337 (0.302, 0.366) |
| QI → U | 0.0624 (0.0269, 0.1020) | 0.408 (0.353, 0.457) |
| I → D | 0.0579 (0.0332, 0.0862) | 0.265 (0.217, 0.318) |
| Q → I | 0.0568 (0.0315, 0.0750) | 0.098 (0.073, 0.121) |
| I → Q | 0.0533 (0.0153, 0.0969) | 0.111 (0.077, 0.156) |
| D → U | 0.0214 (0.0022, 0.0483) | 0.276 (0.178, 0.474) |
| Q → U | 0.0198 (0.0037, 0.0389) | 0.296 (0.209, 0.367) |
| D → I | 0.0180 (0.0092, 0.0275) | 0.155 (0.123, 0.192) |
| D → Q | 0.0177 (0.0058, 0.0315) | 0.184 (0.117, 0.347) |
| U → QI | 0.0097 (0.0022, 0.0181) | 0.371 (0.322, 0.410) |
| I → U | 0.0069 (0.0015, 0.0136) | 0.158 (0.098, 0.223) |
| U → D | 0.0066 (0.0024, 0.0112) | 0.235 (0.176, 0.300) |
| U → Q | 0.0061 (0.0008, 0.0127) | 0.200 (0.119, 0.294) |
| U → I | 0.0037 (0.0009, 0.0071) | 0.147 (0.090, 0.207) |

Note.— The species tree of figure 2 is used, with a single introgression event assumed in each analysis. The full dataset of 1060 loci is analyzed using BPP to estimate the introgression probability ( $\phi$ ) and the introgression time ( $\tau$ ), together with the species divergence times ( $\tau$ s) and population sizes ( $\theta$ s) on the species tree. Introgression events with weak support (U → QI and below) are not considered further.

The time of QI→U introgression was estimated to be 0.000408, very close to the species divergence time at node QIR (0.000417) (fig. 2), suggesting that the introgression was probably a more ancient event. Note that if an introgression event is assigned incorrectly to a daughter branch to the lineage truly involved in introgression, one would expect the estimated introgression time to collapse onto the species divergence time. We thus attempted to place the introgression onto more ancient ancestral branches on the species tree (fig. 2) and finally identified the lineage involved in introgression to be the ancestral species QIRCD. The QIRCD→U introgression had an estimated time that was away from the species divergence times, and the estimated introgression probability (62%) was the highest (table 2).

In the second stage, we added introgression events identified in table 2 onto the binary species tree of figure 2, in the order of their introgression probabilities (table S1). This was applied to the empirical data (split into two halves). Our procedure allows introgression events already in the model to drop out when new introgressions are added to the model. However, this did not happen in the analysis of the *Tamias* dataset. Instead the most important introgression events identified in stage 1 remained to be most important in the joint introgression models constructed in stage 2. Note that multiple introgression events may not be independent. An introgression event significant in stage 1 may not be significant anymore when other introgression events are already included in the model. For example, when the QI→D introgression was already included in the model, none of the introgressions Q→D, D→QI, and I→D were significant. Those introgressions may be expected to lead to similar features in the sequence data, such as reduced sequence divergences between D and Q and between D and I. Similarly, introgression probability for an introgression event often became smaller when other introgressions were added in the model. However, the opposite may occur as well, with the introgression probability becoming higher when other introgression events are included in the model. For example, *φ*_*QIRCD*→*U*_ was estimated to be 54-63% when this was the only introgression assumed in the model, but increased to 60-70% when other introgression events were added in the model (table S1).

The results for the two halves of the empirical data were largely consistent, especially concerning the most important introgression events with high introgression probabilities. We thus arrived at a joint introgression model with five unidirectional introgression events and one bidirectional introgression event (table S1, fig. 3).

### Estimation of introgression probabilities and species divergence/introgression times

Finally, we fitted the joint introgression model of figure 3 to the full data of 1060 loci, as well as the two halves. Estimates of introgression probabilities and introgression times as well as other parameters (*θ* s and *τ*s) under the joint model are given in table 3. This model is very parameter-rich. While the divergence/introgression times are well estimated, the population sizes for ancestral species are poorly estimated, especially for those populations with a very short time duration. The current version of BPP assigns a different *θ* parameter for the same population before and after each introgression event (for example, branch Q in figure 3 is broken into four segments by three introgression events (Q↔I, D→Q, and Q→U) and are assigned four independent population size parameters.

**Table 3.** Posterior means and 95% HPD CIs (in parentheses) of parameters under the MSci model of figure 3 obtained from BPP analyses of three real datasets (the two halves and the full dataset) and a simulated dataset.

| Parameters | First half, 530 loci | Second half, 530 loci | Full data, 1060 loci | Simulation, 1060 loci |
| --- | --- | --- | --- | --- |
| Population sizes θs ($\times 10^{-3}$) | | | | |
| $\theta_Q$ | 1.080 (0.747, 1.450) | 1.024 (0.523, 1.539) | 1.087 (0.767, 1.416) | 1.037 (0.790, 1.518) |
| $\theta_I$ | 1.612 (0.979, 2.374) | 1.973 (0.815, 3.361) | 2.073 (1.121, 2.999) | 1.947 (1.136, 3.129) |
| $\theta_R$ | 0.345 (0.279, 0.408) | 0.381 (0.315, 0.443) | 0.349 (0.305, 0.392) | 0.347 (0.310, 0.392) |
| $\theta_C$ | 0.469 (0.389, 0.551) | 0.469 (0.395, 0.538) | 0.475 (0.420, 0.529) | 0.481 (0.424, 0.539) |
| $\theta_D$ | 3.730 (2.252, 5.802) | 5.199 (2.939, 8.032) | 5.012 (2.885, 7.774) | 4.673 (2.547, 7.049) |
| $\theta_U$ | 1.205 (0.995, 1.418) | 1.265 (1.044, 1.493) | 1.234 (1.087, 1.383) | 1.236 (1.092, 1.365) |
| $\theta_S$ | 0.791 (0.684, 0.907) | 0.938 (0.814, 1.058) | 0.869 (0.786, 0.951) | 0.830 (0.746, 0.913) |
| $\theta_{QIRCDUS}$ | 11.02 (9.454, 12.58) | 10.64 (9.166, 12.15) | 10.80 (9.657, 11.93) | 11.05 (9.979, 12.15) |
| $\theta_{QIRCDU}$ | 0.621 (0.295, 0.986) | 0.624 (0.303, 0.958) | 0.645 (0.373, 0.925) | 0.614 (0.342, 0.913) |
| $\theta_{QIRCD}$ | 1.591 (0.236, 3.750) | 1.542 (0.214, 3.668) | 1.984 (0.249, 4.726) | 1.849 (0.243, 4.503) |
| $\theta_{QIRC}$ | 3.214 (0.511, 6.249) | 1.422 (0.239, 3.191) | 2.322 (0.354, 4.697) | 2.512 (0.382, 4.956) |
| $\theta_{QIR}$ | 2.372 (0.491, 4.284) | 1.633 (0.609, 2.734) | 2.218 (1.104, 3.372) | 2.143 (0.987, 3.018) |
| $\theta_{QI}$ | 0.915 (0.184, 2.144) | 0.839 (0.174, 1.982) | 0.594 (0.170, 1.233) | 0.622 (0.190, 1.337) |
| $\theta_J$ | 0.970 (0.742, 1.218) | 1.274 (1.002, 1.545) | 1.126 (0.947, 1.308) | 1.089 (0.878, 1.316) |
| $\theta_K$ | 0.938 (0.180, 2.231) | 1.061 (0.181, 2.554) | 1.267 (0.184, 2.456) | 1.017 (0.178, 2.395) |
| $\theta_L$ | 1.503 (0.240, 3.374) | 1.062 (0.228, 2.214) | 1.288 (0.240, 2.938) | 1.416 (0.213, 3.476) |
| $\theta_M$ | 0.387 (0.238, 0.542) | 0.442 (0.270, 0.614) | 0.364 (0.239, 0.493) | 0.329 (0.191, 0.428) |
| $\theta_N$ | 0.938 (0.143, 2.338) | 0.814 (0.149, 2.080) | 0.906 (0.171, 2.212) | 0.864 (0.174, 2.213) |
| $\theta_O$ | 0.591 (0.432, 0.757) | 0.528 (0.389, 0.676) | 0.536 (0.429, 0.645) | 0.520 (0.380, 0.663) |
| $\theta_P$ | 3.975 (0.431, 9.050) | 2.693 (1.354, 4.363) | 3.200 (1.851, 5.091) | 3.641 (1.873, 5.624) |
| $\theta_T$ | 0.531 (0.146, 1.317) | 0.409 (0.183, 0.752) | 0.368 (0.243, 0.481) | 0.469 (0.275, 0.654) |
| $\theta_V$ | 1.574 (0.294, 3.570) | 1.867 (0.330, 4.556) | 1.851 (0.348, 3.861) | 1.894 (0.400, 3.907) |
| $\theta_W$ | 0.414 (0.206, 0.634) | 0.437 (0.273, 0.589) | 0.427 (0.326, 0.507) | 0.491 (0.380, 0.597) |
| $\theta_X$ | 0.648 (0.146, 1.651) | 2.052 (0.412, 5.049) | 1.194 (0.281, 2.437) | 1.231 (0.410, 2.368) |
| $\theta_Y$ | 1.021 (0.220, 2.547) | 1.332 (0.367, 2.657) | 1.555 (0.240, 3.733) | 1.496 (0.292, 3.482) |
| Speciation/Hybridization times τs ($\times 10^{-3}$) | | | | |
| $\tau_{QIRCDUS}$ | 3.284 (2.775, 3.805) | 3.665 (3.164, 4.173) | 3.518 (3.124, 3.919) | 3.691 (3.265, 3.944) |
| $\tau_{QIRCDU}$ | 2.088 (1.557, 2.655) | 2.259 (1.793, 2.744) | 2.151 (1.753, 2.547) | 2.238 (1.775, 2.717) |
| $\tau_{QIRCD}$ | 0.775 (0.657, 0.880) | 0.741 (0.652, 0.823) | 0.740 (0.653, 0.819) | 0.775 (0.661, 0.873) |
| $\tau_{QIRC}$ | 0.566 (0.450, 0.681) | 0.653 (0.554, 0.744) | 0.616 (0.533, 0.699) | 0.589 (0.497, 0.677) |
| $\tau_{QIR}$ | 0.387 (0.315, 0.464) | 0.442 (0.371, 0.513) | 0.395 (0.340, 0.447) | 0.372 (0.328, 0.414) |
| $\tau_{QI}$ | 0.309 (0.262, 0.358) | 0.352 (0.295, 0.408) | 0.328 (0.291, 0.365) | 0.335 (0.301, 0.370) |
| $\tau_J = \tau_K = \tau_{QIRCD \rightarrow U}$ | 0.876 (0.756, 0.996) | 0.804 (0.702, 0.907) | 0.820 (0.739, 0.904) | 0.823 (0.758, 0.936) |
| $\tau_L = \tau_M = \tau_{QI \rightarrow D}$ | 0.328 (0.273, 0.378) | 0.376 (0.313, 0.443) | 0.352 (0.311, 0.392) | 0.360 (0.324, 0.393) |
| $\tau_N = \tau_O = \tau_{Q \rightarrow U}$ | 0.278 (0.205, 0.345) | 0.299 (0.194, 0.379) | 0.303 (0.249, 0.352) | 0.333 (0.257, 0.397) |
| $\tau_P = \tau_T = \tau_{D \rightarrow Q}$ | 0.178 (0.088, 0.289) | 0.125 (0.074, 0.173) | 0.114 (0.081, 0.149) | 0.119 (0.072, 0.172) |
| $\tau_V = \tau_W = \tau_{D \rightarrow I}$ | 0.126 (0.053, 0.244) | 0.095 (0.047, 0.140) | 0.073 (0.055, 0.113) | 0.082 (0.053, 0.112) |
| $\tau_X = \tau_Y = \tau_{Q \leftrightarrow I}$ | 0.079 (0.041, 0.117) | 0.035 (0.017, 0.064) | 0.055 (0.031, 0.069) | 0.054 (0.027, 0.087) |
| Introgression probabilities ϕs | | | | |
| $\phi_{QIRCD \rightarrow U}$ | 0.672 (0.497, 0.845) | 0.660 (0.519, 0.797) | 0.653 (0.536, 0.769) | 0.650 (0.534, 0.769) |
| $\phi_{QI \rightarrow D}$ | 0.130 (0.082, 0.180) | 0.095 (0.044, 0.150) | 0.119 (0.078, 0.161) | 0.125 (0.087, 0.174) |
| $\phi_{Q \rightarrow U}$ | 0.031 (0.007, 0.056) | 0.009 (0.000, 0.025) | 0.018 (0.003, 0.034) | 0.023 (0.005, 0.040) |
| $\phi_{D \rightarrow Q}$ | 0.030 (0.000, 0.088) | 0.020 (0.004, 0.038) | 0.013 (0.005, 0.022) | 0.015 (0.005, 0.029) |
| $\phi_{D \rightarrow I}$ | 0.007 (0.000, 0.019) | 0.013 (0.003, 0.026) | 0.008 (0.003, 0.015) | 0.009 (0.003, 0.016) |
| $\phi_{Q \leftrightarrow I}$ | 0.030 (0.010, 0.056) | 0.036 (0.018, 0.056) | 0.029 (0.018, 0.041) | 0.029 (0.014, 0.046) |
|  | 0.026 (0.006, 0.051) | 0.010 (0.001, 0.019) | 0.015 (0.006, 0.025) | 0.020 (0.007, 0.032) |

Note.— The two introgression probabilities for the bidirectional introgression event between Q and I are $\phi_{Q \rightarrow I}$ (above) and $\phi_{I \rightarrow Q}$ (below).

Two ancient introgression events, QIRCD→U and QI→D, had the highest introgression probabilities, estimated from the full data at 65% and 12%, respectively (table 3).

While estimates of times for the recent divergences (*τ*_*QI*_, *τ*_*QIR*_, *τ*_*QIRC*_, *τ*_*QIRCD*_) are nearly identical between the MSC model ignoring gene flow and the MSci model incorporating gene flow, the age of the *T. quadrivittatus* clade (node QIRCDU) was drastically different between the two models (fig. 4). This is clearly due to the fact that the MSC model ignores the introgression QIRCD→U introgression, which had introgression probability 65%. Ignoring gene flow when it exists is expected to lead to underestimation of species divergence times (Leaché *et al*., 2014). Note that in the MSC model the sequence divergence time has to be larger than the species divergence time (*t*_*XY*_ *> τ*_*XY*_ for any pair of species *X* and *Y*). The estimate of species divergence time (*τ*) is influenced more by the minimum sequence divergence than by the average sequence divergence. If gene flow is present between species and is ignored in the MSC model, the reduced sequence divergence resulting from gene flow will be misinterpreted as a more recent species divergence.

**Figure 4:**
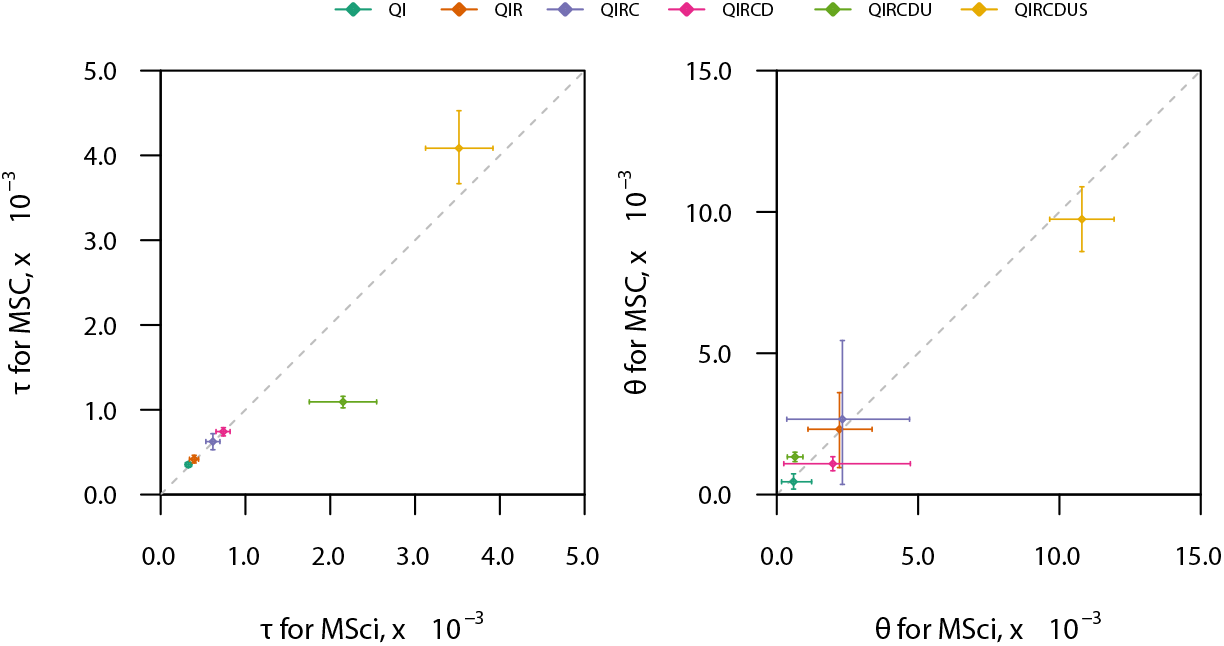
Scatterplot of posterior means and 95% HPD CIs for the six species divergence times (*τ*s) and six ancestral population sizes (*θ*s) obtained from BPP analyses of the full data of 1060 loci under the MSC model ignoring gene flow (fig. 2) and under the MSci model with introgression (fig. 3).

If we used the minimum divergence time of 7 Ma for the outgroup species *T. striatus*, based on Late Miocene date of Dalquest *et al*. (1996), to rescale the estimates of *τ*s under the MSC model without gene flow (fig. 2), we obtained the minimum age for the *T. quadrivittatus* clade to be 1.9 Ma (with 95% HPD CI to be 1.8– 2.0). Here the CI accommodated the uncertainty due to finite amounts of sequence data but not uncertainties in the fossil calibration. Using the same calibration and analyzing four nuclear genes, Sullivan *et al*. (2014, fig. 1) dated the *T. quadrivittatus* clade to 1.8Ma from a maximum likelihood analysis of concatenated data, and to 1.2 Ma (with 95% CI 0.6–2.2) using *BEAST (Heled and Drummond, 2010). Concatenated analysis is known to be biased as it does not accommodate the coalescent process or the stochastic fluctuation of gene tree topologies and divergence times (Ogilvie *et al*., 2017). The *BEAST estimates are comparable to our estimates under the MSC (fig. 2). We then applied the same calibration to rescale the estimates of *τ*s under the MSci model accommodating cross-species gene flow, and obtained the minimum age for the *T. quadrivittatus* clade to be 4.3 Ma (with CI be 3.5–5.1) (fig. 3), much older than the estimates under MSC ignoring gene flow.

### Simulation results

While the analyses of nuclear data by Sarver *et al*. (2021) using HyDe detected no signal for gene flow at all, our BPP analyses of the same data revealed strong evidence of multiple introgression events, involving both sister and non-sister species (fig. 3). We conducted two sets of simulations to examine whether the opposing results may be explained by different efficiency or power of the methods used. We note that the QIRCD→U introgression, the strongest according to our analysis with the introgression probability estimated at 65%, is not detectable as HyDe is unable to detect gene flow between sister species. The next strongest signal is for the QI→D introgression, with the introgression probability ∼12% (fig. 3).

In the first set of simulation, we generated datasets under the joint MSci model of figure 3 for all seven species, but analyzed only data for the quartet Q, R, D and S, focusing on the Q→D introgression. The same datasets were used in the Hyde and BPP analyses, with S used as the outgroup. Note that here the model is misspecified as the data were simulated under the model of figure 3. First, some species were not sampled, and gene flow involved both sampled and unsampled species. Second, some branches on the big species tree merged into one branch on the small tree, so that the population size parameter for the new model is an average of multiple population size parameters in the original large tree.

The Hyde test, with Q specified as the ‘hybrid’ species, detected gene flow at the 1% significance level in 0%, 0%, 0%, 1%, and 11% of replicates, in datasets of 100, 200, 400, 800, and 1600 loci, respectively, so the power was above the nominal 1% only when *L* ≥ 1600 (table 4). Note that a random test that does not use the data should reject the null hypothesis 1% of the time, so that HyDe is conservative. At the size of the *Tamias* dataset (with 1060 loci), HyDe is expected to have little power of detecting even the strongest signal of gene flow identified in our analyses. Estimates of introgression probability by HyDe were often outside the (0, 1) range and were biased (table 4). We also ran Hyde with D specified as the ‘hybrid’ species. The performance was even poorer: the power of detecting gene flow was 0% at all datasizes, and the average estimates of the introgression probability were ∼ 0.52.

**Table 4.** Power of BPP and HyDe tests of Q→D gene flow (fig. 5) and average estimates of introgression probability in 100 simulated replicates.

| # loci | BPP |  | HYDE |  |  |  |
| --- | --- | --- | --- | --- | --- | --- |
| | Power ($\alpha = 1\%$) | Mean $\hat{\phi}$ | Power ($\alpha = 1\%$) | Power ($\alpha = 5\%$) | Mean $\hat{\phi}$ | Proportion of valid estimates |
| 100 | 56% | 0.354 | 0% | 0% | 0.387 | 63% |
| 200 | 32% | 0.218 | 0% | 0% | 0.269 | 62% |
| 400 | 50% | 0.118 | 0% | 2% | 0.236 | 62% |
| 800 | 80% | 0.104 | 1% | 1% | 0.160 | 59% |
| 1600 | 98% | 0.112 | 11% | 13% | 0.142 | 68% |

Note.— The Bayesian test by BPP is considered significant if the posterior mean $\hat{\phi} > 0.01$ and the 95% HPD CI excludes 0.001. HYDE test is conducted at the 1% and 5% levels. In some replicates HYDE produced invalid estimates of the introgression probability, outside the range (0, 1), and only the valid estimates are used to calculate the means. Data were simulated under the joint model of figure 3 and samples from the quartet Q, R, D, and S (outgroup) were analyzed using the species tree of figure 5 to test for gene flow from Q→D. In the HYDE test, Q was specified as the ‘hybrid’ species. See legend to figure 6.

We ran BPP under the MSci model of figure 5 for four species with one introgression event from Q→D. When the same cutoff was used as in the analysis of the real data (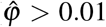 and 95% HPD CI excluding 0.001), gene flow was detected in 56%, 32%, 50%, 80%, and 98% of replicates, in datasets of 100, 200, 400, 800, and 1600 loci, respectively. Thus the BPP analysis was more powerful than the HyDe test, and had high chances of detecting gene flow at the datasize of the *Tamias* dataset.

**Figure 5:**
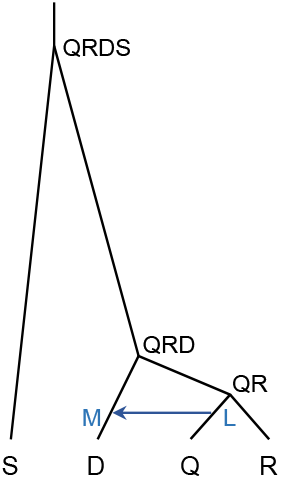
Species tree for a subset of four species (Q, R, D, and S) based on the tree of figure 3, with one introgression event from Q→D, used in simulation to examine the power of HyDe and BPP to detect gene flow.

While HyDe can estimate only two parameters from the site-pattern counts (the internal branch length in coalescent units on the species tree and the introgression probability), BPP analysis of the same data estimates all 14 parameters in the MSci model (fig. 5): 4 species divergence/introgression times (*τ*s), 9 population sizes (*θ* s), and one introgression probability (*φ*_*Q*→*D*_). The parameter estimates are summarized in figure 6. While the model is misspecified, the four divergence/introgression times had the same definition as in the original correct model (fig. 3); these were all very well estimated with small CIs. The introgression probability *φ*_*Q*→*D*_ was accurately estimated with narrow 95% HPD CIs when ≥ 800 loci were used. Population size parameters for short branches were poorly estimated due to lack of coalescent events in those populations. Note that sampling only four species means that multiple branches in the original species tree (fig. 3) are merged into one branch in the quartet tree. The population size for the branch in the quartet tree may thus be a kind of average of the population sizes in the different populations represented by the merged branch. Here we consider two averages, the arithmetic mean *A* and the harmonic mean *H*:

**Figure 6:**
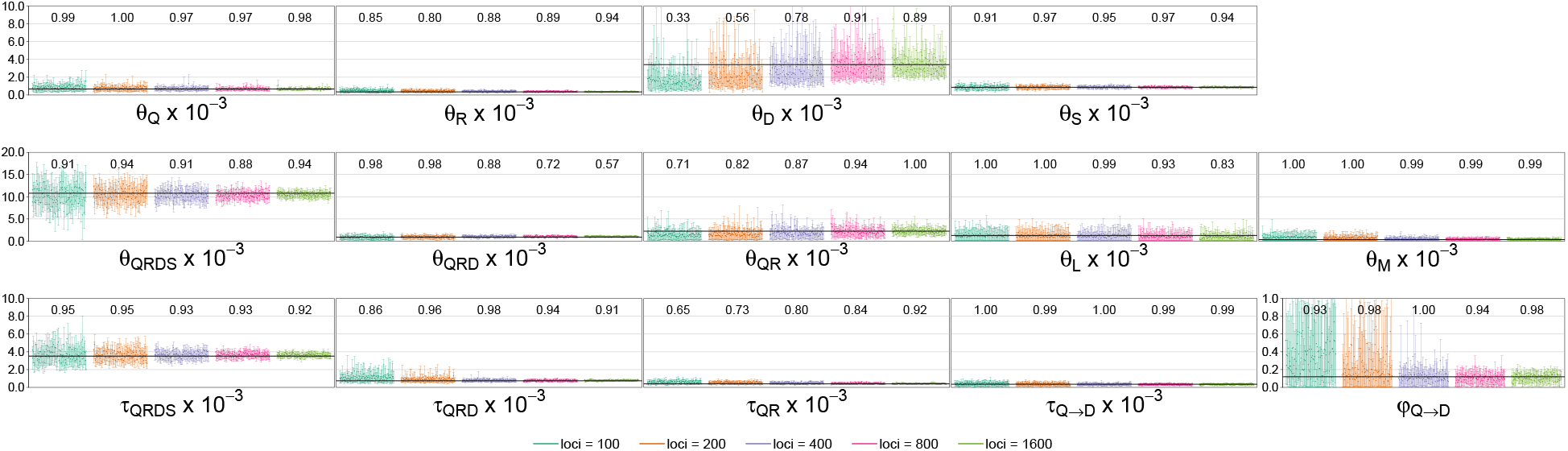
Posterior means and 95% HPD CIs for parameters in the MSci model of figure 5 in BPP analyses of 100 replicate datasets at different datasizes (with the number of loci to be = 100, 200, 400, 800, 1600). Multilocus sequence alignments were simulated under the joint MSci model of figure 3 with seven species and six introgression events (five unidirectional and one bidirectional introgressions), but only samples from the quartet species Q, R, D, and S (outgroup) were analyzed assuming the MSci model of figure 5 with one introgression event from Q→D. The probability that the CI includes the true value (coverage probability) is shown above the CI bars. For population size parameters (*θ* s) that correspond to multiple populations on the true tree of figure 3, the true value is calculated as the arithmetic mean (eq. 1). For example branch QRD in figure 5 corresponds to three branches in figure 3.

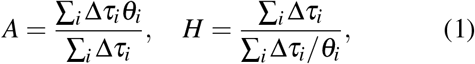

where *θ*_*i*_ is the *θ* value for time segment *i*, and Δ*τ*_*i*_ is the time duration of the segment. For example, branch QRD in figure 5 corresponds to three branches in figure 3, and the weighted means of parameter estimates (*θ*_*QIRCD*_ = 0.00198, *θ*_*J*_ = 0.00113, and *θ*_*QIRCDU*_ = 0.00065) (table 3) gave *A* = 0.00091 and *H* = 0.00083.The arithmetic mean is used to represent the true value in figure 3.

In the second set of simulations, we used the joint MSci model of figure 3 to generate two datasets of 1060 loci, just like the real dataset, to confirm that parameter estimates from datasets of this size using BPP achieve good accuracy and precision. The estimates from the two datasets were similar, and those for one set are shown in table 3. The posterior means were close to the true values, and the CIs were also similar to those calculated from the real data. Similarly to analyses of the real data, divergence times and population sizes for modern species were well estimated, but ancestral population sizes, in particular those for populations of short time duration, were more poorly estimated.

## DISCUSSION

### Identifiability issue in MSci models

The bidirectional introgression event between *T. quadrivittatus* and *T. cinereicollis* in the joint introgression model (fig. 3) poses an unidentifiability issue (Flouri *et al*., 2020). Two sets of parameter values, Θ and Θ′, with 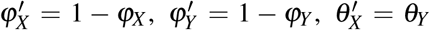, and 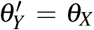, predict identical probability distributions for gene trees or multilocus sequence data and thus cannot be distinguished by such data (fig. 7). The posterior distribution of the parameters will then be bimodal, with the two modes forming perfect mirror images. This is a label-switching type of parameter unidentifiability (Flouri *et al*., 2020; Yang and Flouri, 2021). In our replicate MCMC analyses, the Markov chain visited different modes in different runs, depending on the initial conditions. None of the runs visited both modes, presumably because the dataset of 1060 loci is large so that the mirror modes in the posterior are highly concentrated and well separated. As recommended by Yang and Flouri (2021), in such cases, we summarized samples from the two different modes separately. While the genetic data cannot distinguish the two sets of parameter values, we prefer Θ with small introgression probabilities (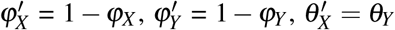 and 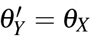), since the alternative, with 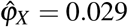 and 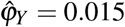, posits mutual near complete replacements, which does not appear plausible (fig. 7).

**Figure 7:**
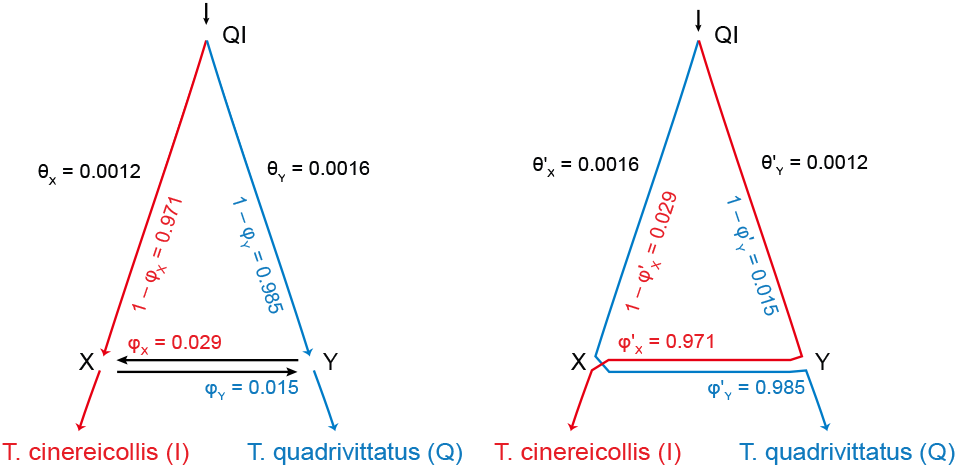
The bidirectional introgression (BDI) event between *T. quadrivittatus* and *T. cinereicollis* in the MSci model of figure 3 poses an unidentifiability issue. Two sets of parameters Θ (left) and Θ′ (right), with 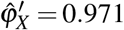 and 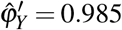, are indistinguishable. The model tree of figure 3 involves seven species and here we show only the part involving the two species. Blue and red lines indicate major routes taken by samples from *T. quadrivittatus* and *T. cinereicollis*, respectively, when one traces the genealogical history of the samples backwards in time.

### The power of heuristic and likelihood methods to detect introgression

HyDe (Blischak *et al*., 2018) is a computationally efficient approach to testing for hybridization/gene flow between two nonsister species, using an outgroup. Suppose there are three populations *P*_1_, *P*_2_, and *P*_3_, related by the phylogeny ((*P*_1_, *P*_2_), *P*_3_). With the outgroup *O*, the parsimony-informative site pattern *ii j j* (where *i* and *j* are any two distinct nucleotides and *ii j j* means *P*_1_ and *P*_2_ have the same nucleotide while *P*_3_ and *O* have another nucleotide) may be considered to match the species tree, while *i j ji* and *i ji j* are the mismatching patterns. Given the binary species tree with no gene flow, the two mismatching patterns have the same probability, but when there is gene flow between *P*_3_ and *P*_1_ (or *P*_2_), they will have different probabilities. The site-pattern counts can thus be used to test for gene flow, as in the so-called *D*-statistic or *ABBA*-*BABA* test (Patterson *et al*., 2012), in which site patterns *ABBA* and *BABA* correspond to *i j ji* and *i ji j*, respectively. Under certain assumptions, the site pattern counts may also be used to estimate the introgression probability as in Hyde (Blischak *et al*., 2018). Both the *D*-statistic and HyDe require negligible computation, but both use simple summaries of the multilocus sequence data and fail to make use of all information in the data. Here we discuss a few issues with HyDe, which may be contributing factors for the opposite conclusions reached in testing for introgression in the *Tamias* dataset.

First, both the *D*-statistic and HyDe count the site patterns by merging them across loci, so that the the site-pattern counts are genome-wide averages. One of the hallmarks of cross-species gene flow is that it creates genealogical variations across the genome. As a result, there should be important information about gene flow in the variance of site-pattern counts among loci, but this information is ignored by those methods. In other words, the genealogical history is expected to vary among loci under the multispecies coalescent model (Barton, 2006), influenced by parameters such as the species divergence times, ancestral population sizes, and the rate of gene flow. Under the assumption of no recombination among sites of the same sequence, all sites at the same locus share the genealogical history while differences among sites of the same locus reflect the stochastic fluctuations of the mutation process. When the site patterns are merged across loci, those two sources of variation are confounded (Shi and Yang, 2018; Zhu and Yang, 2021). As a result, certain forms of introgression, such as introgression between sister species, are unidentifiable by the *D*-statistic or HyDe (Jiao *et al*., 2021). In the chipmunk dataset, the strongest signal of introgression is from QIRCD→U, with the estimated introgression probability reaching 65%. Such introgression cannot be detected by HyDe.For introgression between nonsister species, HyDe can detect signals of gene flow, but is unable to identify its direction and timing.

Second, the HyDe test is based on a model of hybrid speciation, which, for the model of figure 5, would assume inflow, i.e., gene flow from M to L, rather than outflow from L to M (as in the model of fig 5). Furthermore, a symmetry was assumed in divergence times and population sizes (i.e., *τ*_*M*_ = *τ*_*QR*_ and *θ*_*M*_ = *θ*_*QR*_ in fig. 5) (Blischak *et al*., 2018; Kubatko and Chifman, 2019). Indeed those authors assumed the same population size for all species on the species tree. In other words, when applied to simulated data in this study under the model of figure 5, the HyDe model assumed introgression from M to L (with *T. quadrivittatus* to be the hybrid species) with *τ*_*M*_ = *τ*_*QR*_. Those assumptions were not met in our simulation, which may partly explain why HyDe frequently produced invalid estimates of introgression probability in our simulation (table 4). Our BPP analyses identified multiple introgression events in the *Tamias* group (fig. 3), but none of them represented a case of hybrid speciation. While HyDe was described and used as a well-justified approach for assessing introgression in general (Blischak *et al*., 2018; Kubatko and Chifman, 2019; Sarver *et al*., 2021), we suggest that it is applicable to the case of hybrid speciation only. We note that in simulations where all assumptions of the HyDe model were met, the method produced well-behaved estimates of introgression probabilities (Blischak *et al*., 2018; Flouri *et al*., 2020).

Third, the approaches taken by HyDe to accommodate multiple samples per species and heterozygote sites in diploid genomes are problematic. When multiple samples are available in the species quartet, HyDe counts site patterns in all combinations of the quartet. Let the numbers of sequences for species *P*_1_, *P*_2_, *P*_3_ and *O* be *n*_1_, *n*_2_, *n*_3_, and *n*_*O*_. There are then *n*_1_×*n*_2_×*n*_3_×*n*_*O*_ combinations in which one sequence is sampled per species, and HyDe counts site patterns in all of them. This ignores the lack of independence among the quartets and grossly exaggerates the sample size. At the same time, HyDe does not use the information in the samples from the four species efficiently. For example, multiple samples per species are very informative about the population size for that species, but that information is not used by HyDe, even though the method assumes the same population size for all populations on the species tree. In a full likelihood method such as bpp, all sequences at the same locus, both from the same species and from different species, are related through a gene tree, and genealogical information at the locus is fully used.

Similarly heterozygotes are treated as ambiguities in HyDe. If the site pattern is AGRG, with R representing a A/G heterozygote, HyDe adds 0.5 each to the site patterns *i j j j* (for AGGG) and *i ji j* (for AGAG). The heterozygote R means both A and G rather than an unknown nucleotide that is either A or G. In theory the proportion of heterozygotes in each diploid genome should be a very informative estimate for *θ* for that population, but such information is not used by HyDe. In BPP, heterozygous sites are resolved into their underlying nucleotides using an analytical integration algorithm (Gronau *et al*., 2011; Flouri *et al*., 2018). Uncertainties in the genotypic phase of multiple heterozygous sites in an unphased diploid sequence are accommodated by averaging over all possible heterozygote phase resolutions, weighting them using their relative likelihoods based on the sequence alignment at the locus. Simulations suggest that this approach has nearly identical statistical performance from using fully phased haploid genomic sequences, which could be generated, for example, by costly single-molecule cloning and sequencing (Huang *et al*., 2021).

### Introgression in *T. quadrivittatus* chipmunks

The joint introgression model for *Tamias* (fig. 3) was determined using a stepwise procedure for iteratively adding introgression events to the species tree. This approach may not be feasible for all empirical studies. If the species tree is large and highly uncertain, there may be too many candidate models to evaluate. The *Tamias* data tested here include only six species, and the first stage of our procedure involved 16 possible introgression events, so that it was computationally feasible.

Given the extensive mitochondrial introgression in the *T. quadrivittatus* group of chipmunks (Sullivan *et al*., 2014; Sarver *et al*., 2017, 2021), introgression affecting the nuclear genome was expected (Sarver *et al*., 2021). Thus the failure to detect any significant evidence of nuclear introgressions in the HyDe analysis was surprising. Sarver *et al*. (2021) discussed the accumulating evidence for cytonuclear discordance in the patterns of introgression (Bonnet *et al*., 2017; McElroy *et al*., 2020; Sarver *et al*., 2021), as well as possible roles of purifying selection affecting the coding genes or exons that make up the nuclear dataset being analyzed. Our re-analyses of the same data using BPP suggests that a simpler explanation is that the heuristic methods used by Sarver *et al*. (2021) do not make an efficient use of information in the multilocus sequence alignments and thus lack power: we detect significant and robust evidence of gene flow using BPP, involving both sister species and nonsister species.

Our estimates suggest that species involved in excessive mitochondrial introgression tend to be those involved in nuclear introgression as well. Consistent with the finding that *T. dorsalis* was a universal recipient of mtDNA from other species (Sullivan *et al*., 2014), we detected evidence for multiple introgressions into *T. dorsalis*, with *φ >* 5% (table 2). We also detected evidence for nuclear introgression involving *T. dorsalis* as the donor species: *T. dorsalis* → *T. quadrivittatus, T. dorsalis* → *T. cinereicollis*, and *T. dorsalis* → the *T. quadrivittatus* + *T. cinereicollis* common ancestor.

While the rate of introgression for the nuclear genome may be lower than the rate of mitochondrial introgression, our results suggest overall consistency between the nuclear and mitochondrial genomes in the species involved in introgression in the *T. quadrivittatus* group. It appears beyond doubt that introgression has affected the nuclear as well as the mitochondrial genomes in the group. It will be useful to generate more genomic data, especially the noncoding parts of the nuclear genome, including more species from the genus, to produce more precise estimates of introgression rates. It will also be interesting to examine whether the noncoding and coding regions of the genome give consistent signals concerning species divergences and cross-species gene flow. In a few genomic analyses, of the gibbbons (Shi and Yang, 2018), the *Anopheles gambiae* group of African mosquitoes (Thawornwattana *et al*., 2018), and the *Heliconius* butterflies (Thawornwattana *et al*., 2021), where the coding and noncoding regions of the genome were analzed as separate datasets, the two types of data produced highly consistent results, with the estimated divergence times (*τ*s) and population size parameters (*θ* s) to be nearly perfectly proportional, and with the estimated introgression rates to be very similar, suggesting that the main effect of purifying selection removing deleterious nonsynonymous mutations is to reduce the neutral mutation rate in the coding regions of the genome, but both types of data can be used as effective markers to study the history of species divergence and gene flow.

## ACKNOWLEDGEMENTS

This study has been supported by a Biotechnology and Biological Sciences Research Council grant (BB/T003502/1) and a BBSRC equipment grant (BB/R01356X/1) to Z.Y. and an NSF grant (NSF-SBS-2023723) to A.D.L.

---

**Table S1.** Posterior means and 95% HPD CIs (in parentheses) of introgression probabilities (*φ*) and introgression times (*τ*) in the stepwise construction of the MSci model, applied to datasets of the two halves.

| Model | First half |  | Second half |  |
| --- | --- | --- | --- | --- |
| | $\phi$ | $\tau (\times 10^{-3})$ | $\phi$ | $\tau (\times 10^{-3})$ |
| 1 QIRCD → U | 0.5372 (0.2471, 0.8012) | 0.841 (0.702, 0.980) | 0.6319 (0.3316, 0.8688) | 0.906 (0.740, 1.045) |
| 2 QIRCD → U | 0.6044 (0.3856, 0.8090) | 0.866 (0.769, 0.971) | 0.6895 (0.4893, 0.8726) | 0.917 (0.795, 1.032) |
| QI ↔ D | 0.1339 (0.0888, 0.1800) | 0.341 (0.302, 0.381) | 0.0668 (0.0367, 0.0992) | 0.317 (0.271, 0.358) |
|  | 0.0194 (0.0001, 0.0452) |  | 0.0205 (0.0007, 0.0402) |  |
| 3 QIRCD → U | 0.6008 (0.3565, 0.8250) | 0.863 (0.755, 0.978) | 0.7085 (0.4955, 0.9044) | 0.922 (0.791, 1.050) |
| QI → D | 0.1314 (0.0745, 0.1928) | 0.332 (0.282, 0.379) | 0.0821 (0.0345, 0.1369) | 0.325 (0.261, 0.390) |
| Q → D | 0.0095 (0.0006, 0.0360) | 0.154 (0.059, 0.274) | 0.0094 (0.0006, 0.0222) | 0.119 (0.056, 0.233) |
| 4 QIRCD → U | 0.5930 (0.3616, 0.8094) | 0.856 (0.750, 0.962) | 0.6770 (0.4616, 0.8725) | 0.907 (0.782, 1.029) |
| QI → D | 0.1271 (0.0405, 0.2026) | 0.337 (0.284, 0.399) | 0.0856 (0.0352, 0.1450) | 0.332 (0.273, 0.392) |
| I → D | 0.0170 (0.0009, 0.0918) | 0.144 (0.047, 0.307) | 0.0117 (0.0010, 0.0245) | 0.156 (0.080, 0.221) |
| 5 QIRCD → U | 0.5933 (0.3415, 0.8251) | 0.854 (0.746, 0.975) | 0.7002 (0.5098, 0.8764) | 0.911 (0.794, 1.027) |
| QI → D | 0.1234 (0.0808, 0.1688) | 0.302 (0.257, 0.349) | 0.0963 (0.0601, 0.1339) | 0.328 (0.284, 0.371) |
| Q ↔ I | 0.0403 (0.0127, 0.0722) | 0.111 (0.071, 0.159) | 0.0529 (0.0251, 0.0846) | 0.098 (0.063, 0.135) |
|  | 0.0348 (0.0088, 0.0651) |  | 0.0143 (0.0003, 0.0308) |  |
| 6 QIRCD → U | 0.6565 (0.4766, 0.8306) | 0.840 (0.726, 0.953) | 0.6654 (0.5150, 0.8082) | 0.815 (0.704, 0.927) |
| QI → D | 0.1321 (0.0849, 0.1815) | 0.303 (0.255, 0.353) | 0.1153 (0.0679, 0.1670) | 0.340 (0.291, 0.392) |
| Q ↔ I | 0.0388 (0.0121, 0.0707) | 0.105 (0.055, 0.147) | 0.0490 (0.0222, 0.0785) | 0.088 (0.047, 0.124) |
|  | 0.0320 (0.0074, 0.0596) |  | 0.0134 (0.0003, 0.0284) |  |
| D → U | 0.0115 (0.0001, 0.0243) | 0.265 (0.180, 0.338) | 0.0114 (0.0000, 0.0280) | 0.291 (0.192, 0.365) |
| 7 QIRCD → U | 0.6793 (0.4931, 0.8592) | 0.880 (0.750, 1.009) | 0.6673 (0.5196, 0.8112) | 0.811 (0.708, 0.917) |
| QI → D | 0.1303 (0.0855, 0.1770) | 0.314 (0.259, 0.366) | 0.0969 (0.0596, 0.1358) | 0.330 (0.279, 0.376) |
| Q ↔ I | 0.0369 (0.0108, 0.0682) | 0.096 (0.055, 0.135) | 0.0471 (0.0212, 0.0782) | 0.078 (0.036, 0.127) |
|  | 0.0301 (0.0072, 0.0563) |  | 0.0124 (0.0003, 0.0270) |  |
| Q → U | 0.0293 (0.0068, 0.0537) | 0.257 (0.170, 0.327) | 0.0075 (0.0000, 0.0199) | 0.234 (0.139, 0.327) |
| 8 QIRCD → U | 0.6724 (0.5012, 0.8425) | 0.878 (0.764, 0.990) | 0.6635 (0.5205, 0.8031) | 0.807 (0.705, 0.908) |
| QI → D | 0.1242 (0.0781, 0.1729) | 0.308 (0.259, 0.363) | 0.1079 (0.0591, 0.1610) | 0.354 (0.297, 0.409) |
| Q ↔ I | 0.0365 (0.0126, 0.0644) | 0.098 (0.059, 0.135) | 0.0427 (0.0212, 0.0666) | 0.055 (0.024, 0.091) |
|  | 0.0308 (0.0072, 0.0569) |  | 0.0129 (0.0000, 0.0262) |  |
| Q → U | 0.0279 (0.0061, 0.0515) | 0.252 (0.170, 0.323) | 0.0077 (0.0000, 0.0216) | 0.264 (0.152, 0.356) |
| D → I | 0.0543 (0.0041, 0.1111) | 0.242 (0.123, 0.316) | 0.0158 (0.0032, 0.0299) | 0.125 (0.062, 0.188) |
| 9 QIRCD → U | 0.6719 (0.4971, 0.8448) | 0.876 (0.756, 0.996) | 0.6603 (0.5191, 0.7974) | 0.804 (0.702, 0.907) |
| QI → D | 0.1299 (0.0817, 0.1799) | 0.328 (0.273, 0.378) | 0.0946 (0.0441, 0.1503) | 0.376 (0.313, 0.443) |
| Q ↔ I | 0.0304 (0.0104, 0.0555) | 0.079 (0.041, 0.117) | 0.0358 (0.0180, 0.0562) | 0.035 (0.017, 0.064) |
|  | 0.0264 (0.0063, 0.0506) |  | 0.0096 (0.0010, 0.0193) |  |
| Q → U | 0.0309 (0.0071, 0.0562) | 0.278 (0.205, 0.345) | 0.0093 (0.0000, 0.0250) | 0.299 (0.194, 0.379) |
| D → I | 0.0074 (0.0000, 0.0192) | 0.126 (0.053, 0.244) | 0.0131 (0.0029, 0.0263) | 0.095 (0.047, 0.140) |
| D → Q | 0.0298 (0.0000, 0.0881) | 0.178 (0.088, 0.289) | 0.0200 (0.0039, 0.0384) | 0.125 (0.074, 0.173) |

Note.— Introgression events are added sequentially onto the species tree of figure 2 and those that do not meet our cutoffs ( $\hat{\phi} > 0.01$ and CI excluding 0.001) are grayed out. The final joint introgression model has five unidirectional introgression events and one bidirectional introgression event.

